# Comparative oncology reveals DNMT3B as a molecular vulnerability in soft-tissue sarcoma

**DOI:** 10.1101/2021.12.22.473852

**Authors:** Ashley M. Fuller, Ann DeVine, Ileana Murazzi, Nicola J. Mason, Kristy Weber, T. S. Karin Eisinger-Mathason

## Abstract

Undifferentiated pleomorphic sarcoma (UPS), an aggressive subtype of soft-tissue sarcoma (STS), is exceedingly rare in humans and lacks effective, well-tolerated therapies. In contrast, STS are relatively common in canine companion animals; thus, incorporation of veterinary patients into studies of UPS offers an exciting opportunity to develop novel therapeutic strategies for this rare human disease. Genome-wide studies have demonstrated that UPS is characterized by aberrant patterns of DNA methylation. However, the mechanisms and impact of this epigenetic modification on UPS biology and clinical behavior are poorly understood. Leveraging cell lines and tissue specimens derived from human and canine patients, we discovered that the DNA methyltransferase DNMT3B is overexpressed in UPS relative to normal mesenchymal tissues and associated with a poor prognosis. Consistent with these findings, genetic DNMT3B depletion strongly inhibited UPS cell proliferation and tumor progression. However, existing hypomethylating agents, including the clinically approved drug 5-aza-2’-deoxycytidine and the DNMT3B-inhibiting tool compound nanaomycin A, were ineffective in UPS due to cellular uptake and toxicity issues. Thus, further development of DNMT3B-targeting strategies for these patients is critical.

## INTRODUCTION

Soft tissue sarcomas (STS) are heterogeneous tumors that arise in mesenchymal tissues such as muscle, adipose, and fibrous connective tissue. Although STS account for only ∼1% of all adult cancer cases, greater than 70 histologic subtypes have been identified that differ on the basis of their tissue of origin, molecular features, and clinical behavior (Katz *et al*, 2018). Despite this heterogeneity, the majority of adult STS are karyotypically complex lesions that exhibit a diverse spectrum of mutations and chromosomal abnormalities. As a result, these tumors are typically refractory to treatment with currently available targeted therapies, leaving patients with few options beyond surgical resection or amputation, radiation, and/or chemotherapy (Dufresne *et al*, 2018). Undifferentiated pleomorphic sarcoma (UPS), which predominantly arises in adult skeletal muscle, is a relatively common and particularly aggressive karyotypically complex subtype with a 10-year survival rate of only ∼25% (Fletcher *et al*, 2013; Widemann & Italiano, 2018). Thus, development of more effective and better-tolerated treatment strategies for this disease represents an unmet clinical need.

DNA methylation of tumor suppressor gene promoters and the transcriptional silencing that ensues is a well-established feature of many cancers, including UPS (Cancer Genome Atlas Research Network. Electronic address & Cancer Genome Atlas Research, 2017; Herman & Baylin, 2003; Kawaguchi *et al*, 2006; Koelsche *et al*, 2021; Merritt *et al*, 2018; Renner *et al*, 2013; Seidel *et al*, 2005; Seidel *et al*, 2007; Steele *et al*, 2019). More recent studies have shown that gene body methylation, which is associated with transcriptional upregulation, is an important characteristic of tumor epigenetic landscapes that correlates with increased oncogene dosage (Arechederra *et al*, 2018; Yang *et al*, 2014). In STS, the majority of studies pertaining to aberrant DNA methylation patterns have done little to address the mechanistic basis and functional consequences thereof, focusing instead on improving tumor classification/diagnosis and patient risk stratification (Cancer Genome Atlas Research Network. Electronic address & Cancer Genome Atlas Research, 2017; Kawaguchi *et al*., 2006; Koelsche *et al*., 2021; Merritt *et al*., 2018; Renner *et al*., 2013; Seidel *et al*., 2005; Seidel *et al*., 2007; Steele *et al*., 2019). Therefore, further functional studies in this area, particularly *in vivo*, are necessary. In mammalian cells, DNA methylation is catalyzed by the canonical DNA methyltransferases DNMT1, DNMT3A, and DNMT3B (Lyko, 2018). Thus, these proteins are attractive candidates for enabling a better understanding of the mechanisms and impact of aberrant DNA methylation on UPS biology.

Comparative oncology integrates spontaneous cancers in animals, most notably canine companion animals, into studies of human tumor biology and therapy (Paoloni & Khanna, 2007). In contrast to some murine models of cancer, tumors in pet dogs develop in exclusively immunocompetent settings, exhibit cancer cell-intrinsic and microenvironmental heterogeneity, and can display similar therapeutic responses as their human counterparts (Dennis *et al*, 2011; Gordon *et al*, 2009; Paoloni & Khanna, 2007). Although some highly prevalent human cancers, such as breast, prostate, and lung carcinomas, are rare in dogs (Gordon *et al*., 2009), STS are relatively common, comprising ∼15% of canine malignancies (Rao *et al*, 2020). Moreover, although canine STS are not routinely subjected to detailed subtype classification, several studies have demonstrated that the histologic features of canine sarcomas parallel those of many human subtypes, including UPS (Boerkamp *et al*, 2016; Chijiwa *et al*, 2004; Iwaki *et al*, 2019; Milovancev *et al*, 2015; Schweiger *et al*, 2015). Therefore, canine “models” of STS offer a unique opportunity to advance our understanding of and develop novel therapeutic strategies for this rare human cancer.

Herein, we adopt a comparative oncology approach to probe the functional and clinical significance of canonical DNMTs in UPS. Using cell lines and tissue specimens obtained from both human and canine patients, we demonstrate that DNMT3B overexpression is associated with poor clinical outcomes in UPS, and that functional depletion of this enzyme potently arrests UPS growth both *in vitro* and *in vivo*. We also show that the existing clinically approved DNA hypomethylating agent, 5-aza-2’-deoxycytidine, is an ineffective therapeutic strategy for human and canine UPS patients due to the limited ability of this compound to enter UPS cells/tissues. Similarly, a putative DNMT3B inhibitor, nanaomycin A, manifested *in vivo* toxicity in murine models. Nevertheless, taken together, our data indicate that development of DNMT3B-specific inhibitors for UPS patients is a promising avenue for future research.

## RESULTS

### Therapeutic efficacy of DNA methylation inhibitor 5-aza-2’-deoxycytine in UPS cell lines correlates with expression of nucleoside transporters that enable drug uptake

In clinical oncology settings, inhibition of DNA methylation is achieved with cytidine analogs such as 5-aza-2’-deoxycytidine (Decitabine; DAC). This compound incorporates into DNA and impedes cell proliferation by enabling DNA demethylation and re-expression of previously silenced genes (e.g., cell cycle regulators; pro-apoptotic genes), or by forming irreversible covalent adducts with DNMTs and inducing DNA damage (Derissen *et al*, 2013; Saba, 2007). DAC is FDA-approved for patients with myelodysplastic syndrome and has been studied extensively in *in vitro*, pre-clinical, and clinical studies of hematologic cancers and carcinomas (PubChem: CID 451668, Decitabine; clinicaltrials.gov). However, the therapeutic efficacy of DAC in the context of UPS remains underexplored. To address this knowledge gap, we treated two human patient-derived UPS cell lines with clinically relevant nanomolar doses (Tsai *et al*, 2012) of DAC and assessed its impact on cell proliferation. We also included HT-1080 fibrosarcoma (FS) cells in this analysis because certain FS subtypes, such as myxofibrosarcoma (MFS), are now thought to be genetically indistinguishable from UPS (Cancer Genome Atlas Research Network. Electronic address & Cancer Genome Atlas Research, 2017). DAC significantly reduced the proliferation of HT-1080 and STS-109 cells at one or more time points in a dose-dependent manner (“strong responders”; **Fig 1A**). In contrast, STS-148 cells responded weakly to DAC, with statistically significant, yet biologically modest, reductions in proliferation only apparent after exposure to the highest dose for the longest treatment interval (**Fig 1A**). DAC is taken up into cells via the equilibrative nucleoside transporters hENT1 and hENT2 (encoded by *SLC29A1* and *SLC29A2*, respectively), and *SLC29A1* expression levels inversely correlate with DAC IC_50_ values in human carcinoma and hematologic cancer cell lines (Qin *et al*, 2009). Therefore, we reasoned that differences in *SLC29A1* and/or *SLC29A2* gene expression may underlie the DAC response heterogeneity observed among human UPS and FS cells. Consistent with this hypothesis, the strong responders exhibited significantly greater expression of one (HT-1080) or both (STS-109) nucleoside transporters than the weakly responding STS-148 cells (**Fig 1B**). Thus, we conclude that *SLC29A1* and *SLC29A2* gene expression in human UPS/FS cell lines correlates with DAC responsivity.

**Fig 1.**
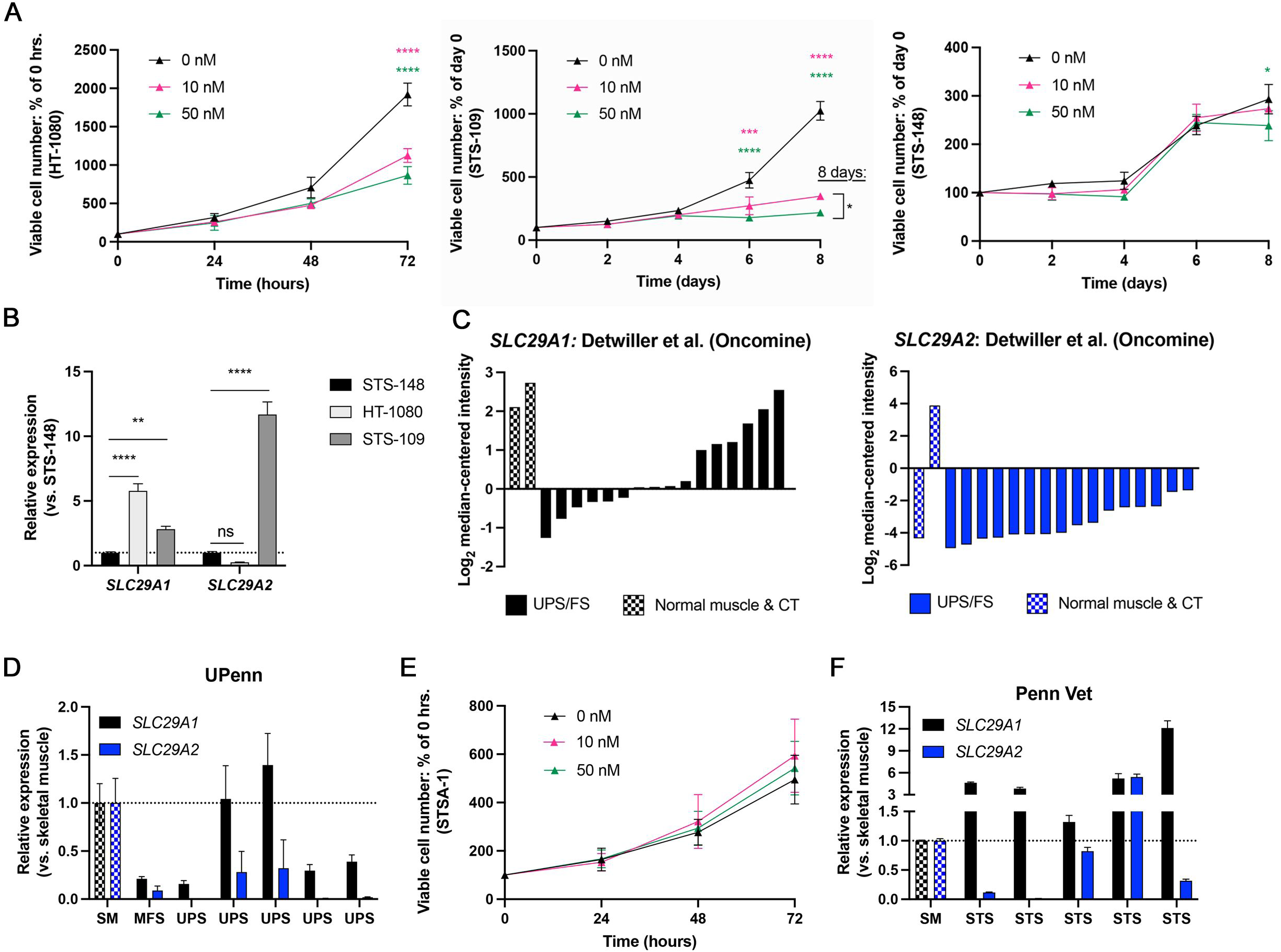
Expression of nucleoside transporter genes *SLC29A1* and *SLC29A2* in human UPS cell lines correlates with 5-aza-2’-deoxycytidine (DAC) responsivity. **A.** Growth curves of human UPS and FS cells treated daily with clinically relevant, nanomolar doses of DAC. Two-way ANOVA with Tukey’s. \**P* < 0.05, \*\*\**P* < 0.001, \*\*\*\**P* < 0.0001. Colored asterisks at each time point indicate significance relative to DMSO. Mean ± SEM. n = 3. **B.** Expression of nucleoside transporter genes in human UPS/FS cell lines. One-way ANOVA with Dunnett’s. \*\**P* < 0.01, \*\*\*\**P* < 0.0001. Mean ± SEM. n = 3. **C.** *SLC29A1* (left) and *SLC29A2* (right) gene expression in UPS and FS tissue specimens from the Detwiller et al. sarcoma dataset (Oncomine). CT = connective tissue. **D.** *SLC29A1* and *SLC29A2* gene expression in human UPS and MFS tumors relative to normal skeletal muscle (SM) tissue (UPenn cohort). Two cDNA samples per specimen. Mean ± SD. **E.** Growth curves of canine STSA-1 cells treated daily with DAC. Two-way ANOVA with Tukey’s. Not significant. Mean ± SEM. n =3. **F.** *SLC29A1* and *SLC29A2* gene expression in canine STS samples relative to normal skeletal muscle (SM) tissue (Penn Vet cohort). Mean ± SD.

To determine the applicability of these relationships to human tissue, we examined *SLC29A1* and *SLC29A2* gene expression patterns in samples from two independent sarcoma patient cohorts: 1) the publicly available Detwiller et al. sarcoma dataset (Detwiller *et al*, 2005; Oncomine) and 2) samples procured through the University of Pennsylvania Surgical Pathology service (“UPenn cohort”). In both cohorts, *SLC29A2* expression was reduced in the majority of UPS/(M)FS relative to normal skeletal muscle and connective tissue specimens, whereas *SLC29A1* expression displayed greater intertumoral heterogeneity (**Fig 1C-D**; **Appendix Table S1**). We also ascertained the relevance of our findings to canine companion animals by leveraging tissue specimens procured by the University of Pennsylvania School of Veterinary Medicine (“Penn Vet cohort”) and canine STSA-1 grade II STS cells (Gentschev *et al*, 2012). The tumor from which these cells were derived was not classified as a particular subtype but resembled UPS with respect to its skeletal muscle origin, pleomorphic histology, and invasive and metastatic behavior (Gentschev *et al*., 2012). Data from canine cells and tissue largely recapitulated that from our human system: STSA-1 cell proliferation was resistant to clinically relevant doses of DAC (**Fig 1E**), similar to human STS-148 cells. Moreover, *SLC29A2* was downregulated in the majority of canine STS specimens relative to normal skeletal muscle (**Fig 1F**), while *SLC29A1* expression tended to be more variable. Taken together, our findings suggest that expression of these transporters in human UPS and canine sarcoma tissue is heterogeneous, as are cellular responses to DAC. Accordingly, we posit that this intertumoral heterogeneity will preclude the clinical efficacy of DAC in many human and canine UPS patients.

### *DNMT3A* and *DNMT3B* regulate UPS cell proliferation *in vitro* and *in vivo*

We next sought to identify more specific and less heterogeneous targets for inhibition of DNA methylation in human UPS and canine sarcoma. We focused on the canonical DNMT proteins DNMT1, DNMT3A, and DNMT3B, which catalyze DNA methylation in mammalian cells (Lyko, 2018). First, we performed cell-based proliferation assays to evaluate the functional role of these enzymes in this tumor context. In addition to our human and canine cell lines, we also used murine KP230 cells, derived from the *Kras*^G12D/+^; *Trp53*^fl/fl^ (KP) genetically engineered mouse model of UPS (Eisinger-Mathason *et al*, 2013). In this model, injection of adeno-Cre virus into the gastrocnemius leads to the development of skeletal muscle tumors that molecularly and histologically recapitulate human UPS (Kirsch *et al*, 2007; Mito *et al*, 2009). Genetic depletion studies in mouse and human cells were performed with gene-specific siRNA pools or lentiviral shRNAs (2 independent shRNAs per target gene). These analyses were not performed in canine STSA-1 cells because rigorously validated canine siRNAs are not widely available, nor were we able to achieve consistent or specific gene silencing with our shRNA lentivirus system (see **Materials and methods**; **Appendix Table S2**). We observed no significant effects on UPS cell growth after depletion of *Dnmt1*/*DNMT1* (**EV Fig. 1A-D**). However, *Dnmt3a*/*DNMT3A* silencing significantly reduced the proliferation of KP230, STS-109, STS-148, and HT-1080 cells (**Fig 2A-D**, **EV Fig 1E-H**), with similar results obtained for *Dnmt3b*/*DNMT3B* knockdown (**Fig 2E-H**, **EV Fig 1I-L**). These data indicate that the KP model successfully recapitulates aspects of human UPS with respect to DNMT function. *DNMT3A* or *DNMT3B* silencing also reduced the number of viable cells over time (relative to the starting number of cells), particularly in STS-109 and STS-148 cells, indicative of cell death (**Fig 2C, G**; **EV Fig 1I**). However, as UPS cell homeostasis in our system is density-dependent, it was unclear whether this phenotype was due to direct sh:*DNMT3A*- or sh:*DNMT3B*-mediated apoptosis, or the inability of these cells to maintain sufficient confluency levels to support viability.

**Fig 2.**
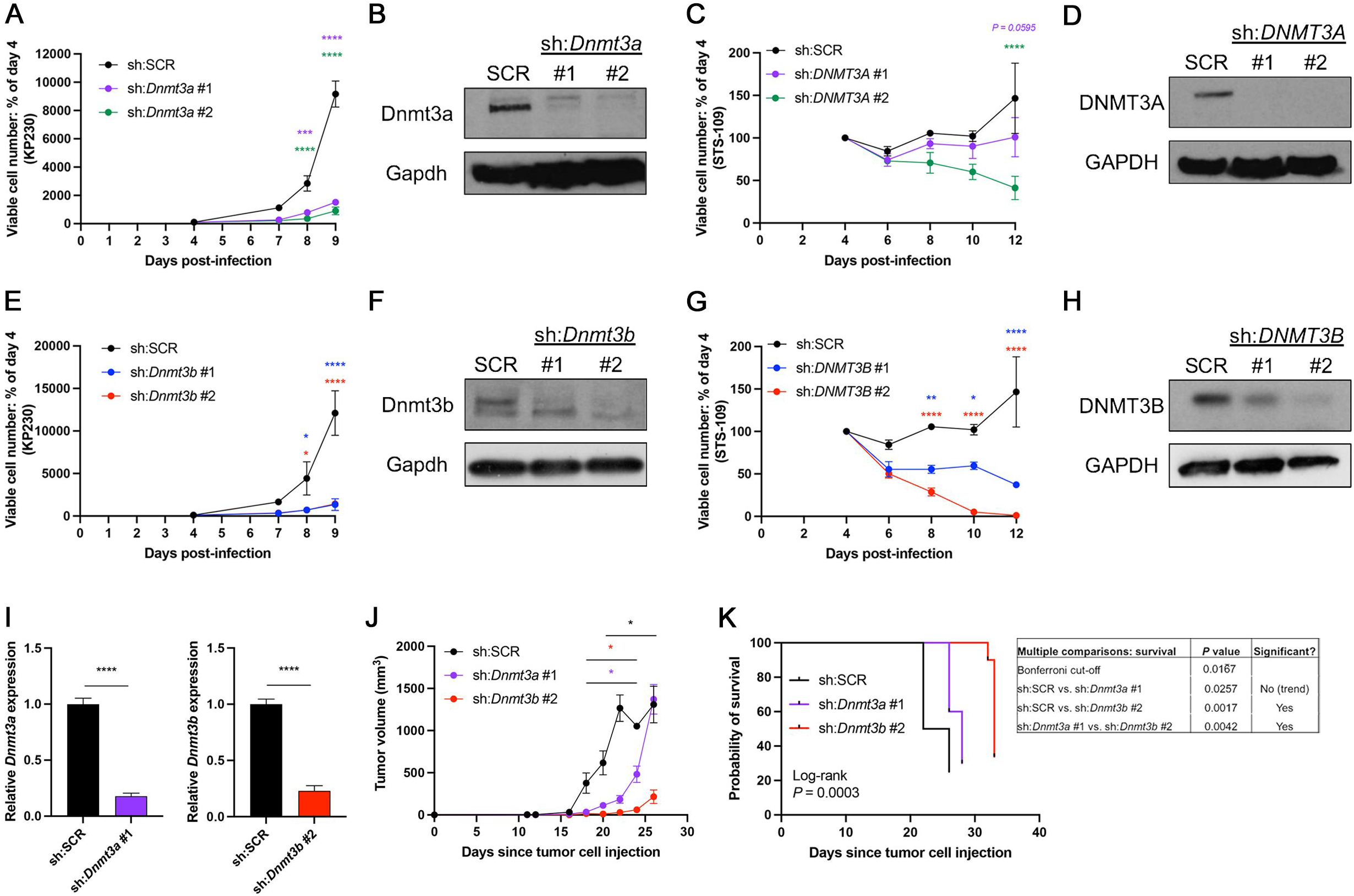
*DNMT3A* or *DNMT3B* depletion inhibits UPS cell proliferation *in vitro* and *in vivo*. **A.** Growth curves of KP230 cells transduced with scrambled control (sh:SCR) or *Dnmt3a*-targeting shRNAs. Two-way ANOVA with Dunnett’s (vs. sh:SCR). \*\*\**P* < 0.001, \*\*\*\**P* < 0.0001; asterisks at each time point indicate significance relative to sh:SCR. Mean ± SEM. n = 3. **B.** Representative Western blot demonstrating *Dnmt3a* knockdown levels in KP230 cells. Image brightness has been enhanced for presentation purposes. **C.** Growth curves of STS-109 cells transduced with control or *DNMT3A*-targeting shRNAs. Two-way ANOVA with Dunnett’s (vs. sh:SCR). \*\*\*\**P* < 0.0001; asterisks at each time point indicate significance relative to sh:SCR. Mean ± SEM. n = 3. **D.** Representative Western blot demonstrating *DNMT3A* knockdown levels in STS-109 cells. Image brightness has been enhanced for presentation purposes. **E.** Growth curves of KP230 cells transduced with control or *Dnmt3b*-targeting shRNAs. Two-way ANOVA with Dunnett’s (vs. sh:SCR). \**P* < 0.05, ****P < 0.0001; asterisks at each time point indicate significance relative to sh:SCR. Mean ± SEM. n = 3. **F.** Representative Western blot demonstrating *Dnmt3b* knockdown levels in KP230 cells. Image brightness and contrast have been enhanced for presentation purposes. The top Dnmt3b band in each lane represents the canonical isoform (isoform 1). **G.** Growth curves of STS-109 cells transduced with control or *DNMT3B*-targeting shRNAs. Two-way ANOVA with Dunnett’s (vs. sh:SCR). \**P* < 0.05, ** *P* < 0.01, ****P < 0.0001; asterisks at each time point indicate significance relative to sh:SCR. Mean ± SEM. n = 3. **H.** Representative Western blot demonstrating *DNMT3B* knockdown levels in STS-109 cells. Image brightness and contrast have been enhanced for presentation purposes. **I.** Gene expression levels of *Dnmt3a* and *Dnmt3b* in KP230 cells orthotopically injected into the gastrocnemius muscles of nude mice. Two-tailed unpaired t-test. \*\*\*\**P* < 0.0001. Mean ± SD. **J.** Nude mouse orthotopic KP230 tumor progression curves. Curves in this panel were terminated when the final mouse in the control group (sh:SCR) was euthanized; however, full tumor growth curves are presented in **EV Fig 1N**. The “dip” in the sh:SCR curve at day 24 occurred because only 1 mouse remained at this time point. Data were analyzed by fitting a mixed model (with Tukey’s multiple comparisons test) due to the presence of missing values. \**P* < 0.05. Colored asterisks indicate significance relative to sh:SCR. Black asterisk indicates significance of sh:*Dnmt3b* vs. sh:*Dnmt3a*. Mean ± SEM. n = 10 tumors (5 mice) per group; 1 tumor in the sh:*Dnmt3b* group did not form. **K.** Survival of orthotopic KP230 tumor-bearing nude mice. Log-rank test with Bonferroni multiple comparisons assessment (table at right). n = 10 tumors (5 mice) per group; 1 tumor in the sh:*Dnmt3b* group did not form.

To determine the role of *Dnmt3a* and *Dnmt3b* in tumor progression, we injected 6-week-old nude mice orthotopically (into the gastrocnemius muscle) with 1 x 10^5^ KP230 cells transduced with a scrambled control (sh:SCR), *Dnmt3a-*, or *Dnmt3b*-targeting shRNA. Each mouse was injected bilaterally to reduce animal use and was euthanized when either intramuscular tumor reached maximum allowed volume (1500 mm^3^). Gene expression levels of *Dnmt3a* and *Dnmt3b* in KP230 cells immediately prior to orthotopic injection, as well as in bulk tumor tissue at the conclusion of the assay, are shown in **Fig 2I** and **EV Fig 1M**, respectively. sh:*Dnmt3b* tumors were significantly smaller than control and/or sh:*Dnmt3a* tumors beginning on day 18 of the study (**Fig 2J**), eventually reaching maximum volume on day 33 (**EV Fig 1N**). sh:*Dnmt3a* tumors were also significantly smaller than control tumors but reached maximum volume more rapidly than sh:*Dnmt3b* tumors. Consistent with this observation, mice bearing sh:*Dnmt3b* tumors experienced significantly longer survival than animals bearing control (*P* = 0.0017) and sh:*Dnmt3a* (*P* = 0.0042) tumors (**Fig 2K**). Animals with sh:*Dnmt3a* tumors also tended to experience prolonged survival compared to control mice, but this comparison did not reach statistical significance. Moreover, among the tumors that had not reached maximum volume at the time of euthanasia, including one sh:*Dnmt3b* tumor that never formed, *Dnmt3b*-deficient tumors were smaller than both control and *Dnmt3a*-deficient tumors (**EV Fig 1O**). Taken together, our results indicate that *Dnmt3a*/*DNMT3A* and *Dnmt3b*/*DNMT3B* both control UPS cell proliferation, but that depletion of *Dnmt3b* more effectively restrains tumor progression *in vivo*.

### *Dnmt3b*/*DNMT3B* deficiency inhibits DNA synthesis in UPS cells

To understand more specifically how *DNMT3B* controls UPS cell proliferation, we performed RNA-seq on human STS-109 cells expressing a scrambled control or one of two independent *DNMT3B*-targeting shRNAs. Cultures were harvested 96 hours after transduction and at 50% confluence to circumvent the effects of density on cell homeostasis and viability. Differential expression analysis was performed to identify genes that were significantly up-or downregulated (FDR *P* < 0.05) in response to both *DNMT3B*-targeting shRNAs compared to the control (**Fig 3A-B**). Metascape analysis of the resulting gene lists indicated that pathways related to autophagy, apoptosis, and protein metabolism were significantly upregulated in sh:*DNMT3B* cells compared to the control (**Fig 3A**), whereas pathways pertaining to DNA replication and cell cycle progression were significantly downregulated (**Fig 3B**). A similar analysis in STS-109 cells expressing *DNMT3A*-specific shRNAs revealed that pathways regulated by *DNMT3A* were distinct from those regulated by *DNMT3B* (**EV Fig 2A-B**). In fact, only 87 genes were significantly upregulated in both sh:*DNMT3A* and sh:*DNMT3B* cells compared to the control, whereas only 65 genes were significantly downregulated (FDR *P* < 0.05; **EV Fig 2C**; **Appendix Table S3**).

**Fig 3.**
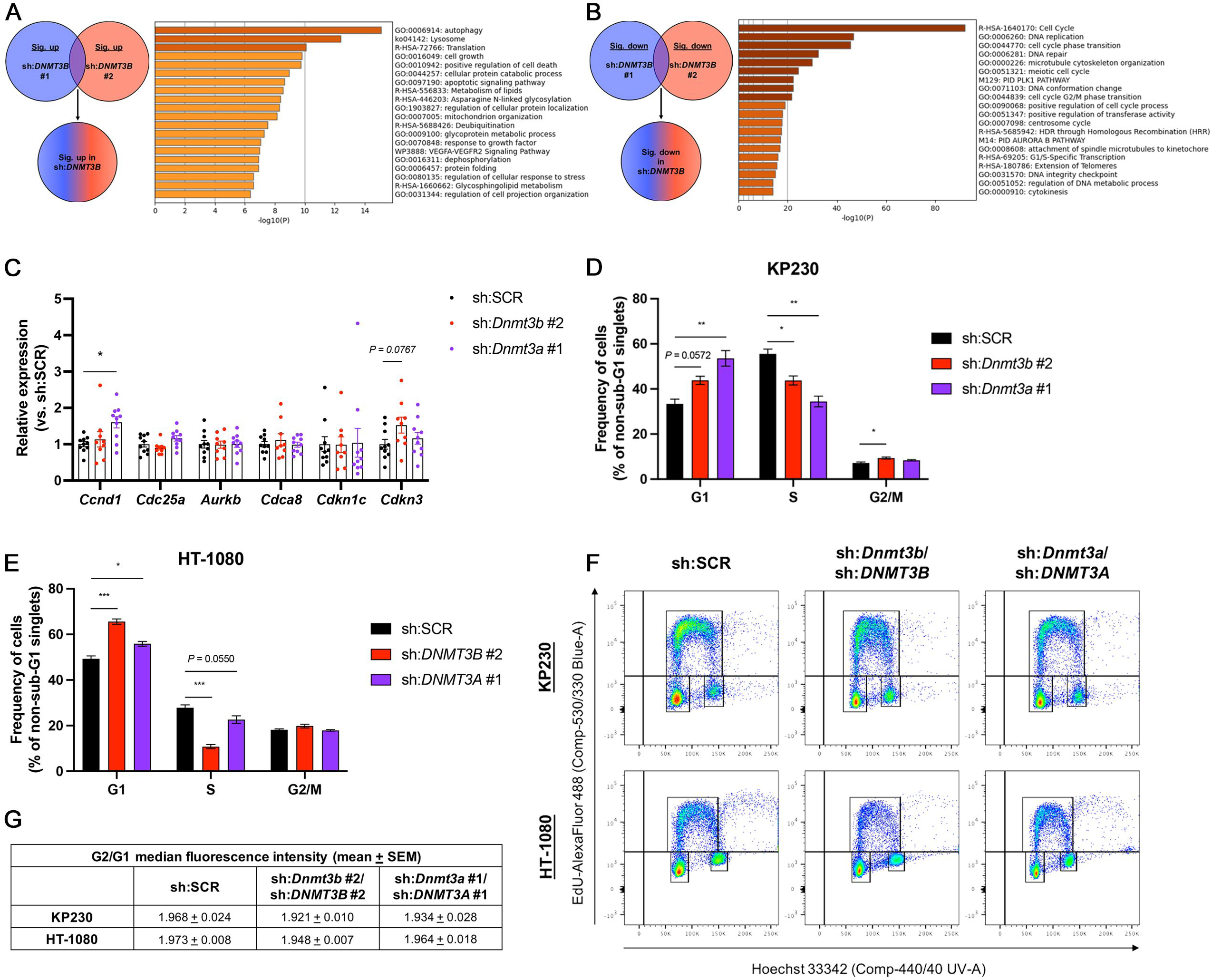
DNMT3B depletion in UPS cells inhibits DNA synthesis. **A.** Schematic and Metascape pathway analysis of genes significantly upregulated in sh:*DNMT3B*-expressing vs. sh:SCR human STS-109 cells (FDR *P* < 0.05). n = 4. **B.** Schematic and Metascape pathway analysis of genes significantly downregulated in sh:*DNMT3B*-expressing vs. sh:SCR human STS-109 cells (FDR *P* < 0.05). n = 4. **C.** Expression of cell cycle-related genes in murine KP230 orthotopic bulk tumor specimens. One-way ANOVA with Dunnett’s (vs. sh:SCR). \**P* < 0.05. Mean ± SEM. n = 10 tumors (5 mice) per group; 1 tumor in the sh:*Dnmt3b* group did not form. **D-E**. Cell cycle phase frequency distributions of KP230 (**D**) and HT-1080 (**E**) cells expressing control, sh:*Dnmt3b*/sh:*DNMT3B*-, or sh:*Dnmt3a*/sh:*DNMT3A*-targeting shRNAs. One-way ANOVA with Dunnett’s (vs. sh:SCR). \**P* < 0.05, \*\**P* < 0.01, \*\*\**P* < 0.001. Mean ± SEM. n = 3. **F.** Representative flow cytometry plots quantified in **D-E** showing the cell cycle phase distributions of control, sh:*Dnmt3b*/sh:*DNMT3B*, and sh:*Dnmt3a*/sh:*DNMT3A* KP230 and HT-1080 cells. **G.** Median fluorescence intensity of Hoechst 33342-DNA staining in the G1- and G2/M-phase cells from panels **D-F**.

We used samples from our murine orthotopic tumor study to explore these potential mechanisms in more detail. We found no evidence of significantly increased apoptosis in sh:*Dnmt3b* vs. sh:SCR specimens (**EV Fig 2D-E**), nor did we observe significant differences in the expression of p62 (**EV Fig 2F-H**), an autophagic substrate that is degraded when autophagy is induced (Mizushima *et al*, 2010; Yoshii & Mizushima, 2017). Therefore, we considered the role of cell cycle regulation. qRT-PCR analysis demonstrated that *Cdkn3*, a cyclin-dependent kinase inhibitor that opposes the G1-S transition (Srinivas *et al*, 2015), was upregulated by ∼50% in sh:*Dnmt3b* compared to sh:SCR tumors (*P =* 0.0767; **Fig 3C**). Due to the heterogeneous nature of bulk tumor RNA, together with the fact that many cell cycle factors are regulated at the protein level, rather than the transcript level (Kronja & Orr-Weaver, 2011; Tanenbaum *et al*, 2015), we performed flow cytometry to obtain a more comprehensive understanding of cell cycle dynamics in *Dnmt3b*/*DNMT3B*-deficient cells (gating strategy provided in **Appendix Fig S1**). In both mouse (KP230) and human (HT-1080) cells, *Dnmt3b*/*DNMT3B* silencing significantly reduced the proportion of cells in S-phase and concomitantly increased the proportion of cells in G1-phase (**Fig 3D-F**). In KP230 cells, there was also a statistically significant, yet modest, increase in the percentage of G2/M-phase cells (**Fig 3D**). Interestingly, similar cell cycle alterations were also observed in sh:*Dnmt3a*/sh:*DNMT3A* cells and bulk tumors (**Fig 3C-F**). For all samples, we confirmed proper gate placement around G1- and G2/M-phase cells using the median fluorescence intensity (MFI) of Hoechst 33342-DNA staining in these populations (**Fig 3G**). All samples had a G2/G1 MFI ratio of ∼2, confirming that the DNA content of G2/M-phase cells was twice that of G1-phase cells. We conclude that *Dnmt3b*/*DNMT3B* depletion impedes UPS cell proliferation by inhibiting DNA synthesis, and that this mechanism also applies in the setting of *Dnmt3a*/*DNMT3A* deficiency.

### DNMT3B inhibitor nanaomycin A potently inhibits UPS cell proliferation *in vitro*

Given the capacity of genetic *Dnmt3b*/*DNMT3B* depletion to restrain UPS growth both *in vitro* and *in vivo*, we sought a pharmacologic approach for clinical translation. Nanaomycin A, a quinone antibiotic, has been described as a DNMT3B-specific inhibitor with tumor-suppressive properties in human colon, lung, and leukemia cell lines (Kuck *et al*, 2010a; Omura *et al*, 1974). Consistent with this report, nanaomycin A exhibited dose-dependent anti-proliferative activity in all five cell lines in our panel, with minimal cytotoxicity (**Fig 4A-C**, **EV Fig 3A-B, Appendix Table S4**). It was most potent in murine KP230 cells, which responded to as little as 10-50 nM after 2-3 days of exposure; no additional benefit was observed after treatment with 100 nM (**Fig 4A, Appendix Table S4**). Growth inhibition in canine and human cells was achieved with slightly higher concentrations (100-500 nM; **Fig 4B-C, EV Fig 3A-B**, **Appendix Table S4**), but all doses used are considered clinically relevant according to commonly accepted preclinical drug screening conventions (Wong *et al*, 2012). We then asked whether nanaomycin A suppresses proliferation via inhibition of DNA synthesis in a manner analogous to *DNMT3B* silencing (gating strategy for STSA-1 cells is shown in **Appendix Fig S2**). Consistent with this hypothesis, nanaomycin A treatment significantly reduced the proportion of human HT-1080 cells in S-phase while concomitantly increasing the percentage of G1-phase cells (**Fig 4D-F**). Murine KP230 and canine STSA-1 cells displayed similar trends, although the observed effects did not reach statistical significance (**EV Fig 3C-E**).

**Fig 4.**
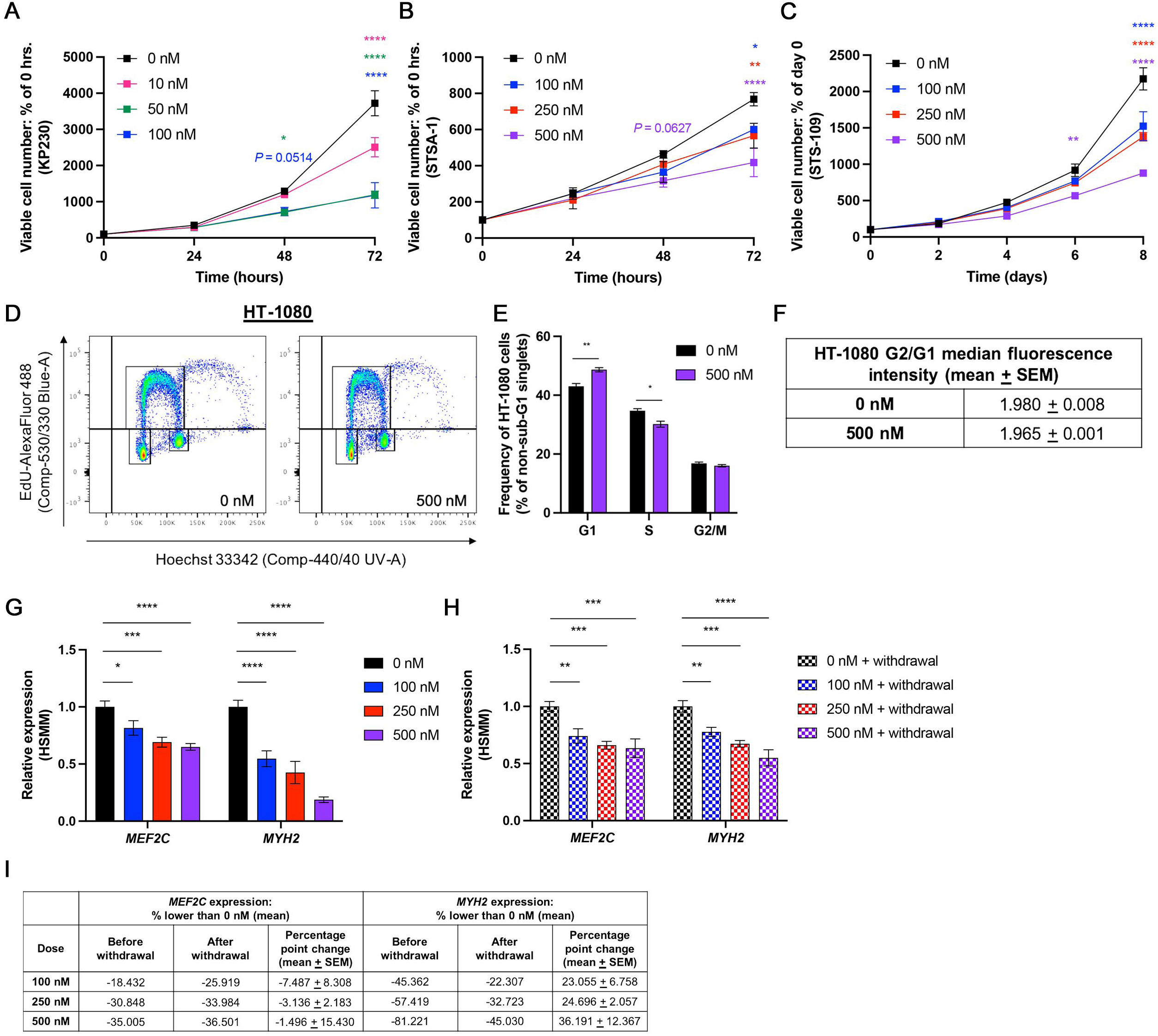
DNMT3B inhibitor nanaomycin a impedes UPS cell proliferation *in vitro*. **A-C.** Growth curves of KP230 (**A**), STSA-1 (**B**), and STS-109 (**C**) cells treated daily with nanaomycin a. Two-way ANOVA with Tukey’s. \**P* < 0.05, \*\**P* < 0.01, \*\*\*\**P* < 0.0001; asterisks represent significance relative to 0 nM. For ease of visualization, <u>only</u> statistical comparisons vs. 0 nM are presented in the figure. Full statistical results for all possible comparisons are shown in **Appendix Table S4**. Mean ± SEM. n ≥ 3. **D**. Representative flow cytometry plots showing the cell cycle phase distributions of the control and nanaomycin a-treated HT-1080 cells quantified in **E** and **F**. **E**. Quantification of HT-1080 cell cycle phase distributions shown in **D**. Two-tailed unpaired t-tests. \**P* < 0.05, \*\**P* < 0.01. Mean ± SEM. n = 3. **F.** Median fluorescence intensity of Hoechst 33342-DNA staining in the G1- and G2/M-phase cells depicted in **D** and **E**. **G.** Expression of human skeletal muscle lineage markers in HSMM myoblasts exposed to nanaomycin a or vehicle control over the course of differentiation to myotubes. One-way ANOVA with Dunnett’s. \**P* < 0.05, \*\*\**P* < 0.001, \*\*\*\**P* < 0.0001. Mean ± SEM. n = 3. **H.** Expression of human skeletal muscle lineage markers in control- or nanaomycin a-treated HSMM myoblasts 5 days after drug withdrawal. \*\**P* < 0.01, \*\*\**P* < 0.001, \*\*\*\**P* < 0.0001. Mean ± SEM. n = 3. **I.** Percentage point change in HSMM lineage marker gene expression after (**H**) vs. before (**G**) nanaomycin a withdrawal.

### Evaluation of nanaomycin A toxicity in normal skeletal muscle cells and murine models

Due to the anti-neoplastic efficacy of nanaomycin A in cell-based models of UPS, we proceeded with *in vitro* functional and toxicity studies of untransformed cells including skeletal muscle myoblasts, widely used *in vitro* models of normal skeletal muscle cells. First, we assessed whether nanaomycin A exposure impairs myoblast differentiation into multi-nucleated myotubes. We treated murine (C2C12), canine (CnSkMC), and human (HSMM) myoblasts with nanaomycin A daily during the course of differentiation, after which expression of skeletal muscle lineage markers (La Rovere *et al*, 2014; Owens *et al*, 2013; Rivera-Reyes *et al*, 2018; Smerdu *et al*, 2005; Strbenc *et al*, 2006) was measured by qRT-PCR (**EV Fig 4A**). Interestingly, we observed distinct species-specific responses. In C2C12s, nanaomycin A only minimally impacted differentiation, significantly reducing the expression of 1 gene, *Myh2*, by ∼20% (**EV Fig 4B**). Conversely, in CnSkMCs, nanaomycin A had inconsistent effects on gene expression at the lowest dose (100 nM), potentially due to mild toxicity, and was overtly cytotoxic at higher doses (≥ 50% and ∼100% cell death at 250 and 500 nM, respectively; **EV Fig 4C**). Yet another distinct pattern was observed for HSMMs, in that differentiation marker expression was uniformly reduced in a dose-dependent manner (**Fig 4G**). Withdrawal of the drug for 5 days did not restore *MEF2C* expression, but did enable recovery of *MYH2* gene expression by up to ∼36 percentage points relative to baseline (**Fig 4H-I**). These data indicate that nanaomycin A impedes human myoblast differentiation, but that this effect is partially reversible when the drug is withdrawn. In addition to testing the effects of nanaomycin A on myoblast differentiation, we also determined whether the compound was cytotoxic to differentiated myotubes. Nanaomycin A had inconsistent effects on the number of viable CnSkMCs, reducing viability in a dose-dependent manner in some experimental replicates, but causing rapid, uniform cell death in others (**EV Fig 4D**). However, it did not impact the viability of C2C12s or HSMMs at any concentration (**EV Fig 4D**).

Largely encouraged by these findings, we evaluated the therapeutic potential of nanaomycin A *in vivo*. We injected 6-week-old nude mice orthotopically with 1 x 10^5^ KP230 cells and administered nanaomycin A or vehicle control i.p. once tumors became palpable. Thereafter, animals were treated every 48 hours for three weeks or until tumors reached maximum allowed volume (1500 mm^3^). Mice treated with 15 mg/kg nanaomycin A exhibited significantly longer survival than mice in the vehicle control group, with two animals in the 15 mg/kg group experiencing near-complete cessation of tumor growth (**EV Fig 4E-F**). Unfortunately, however, these same two mice became cachectic and were humanely euthanized prior to the study endpoint. Similarly, body weights of the remaining mice in the 15 mg/kg group were significantly lower than those of control mice (**EV Fig 4G**). In contrast, mice treated with 7.5 mg/kg nanaomycin A did not show signs of overt toxicity; however, this dose did not significantly extend survival relative to the vehicle control (**EV Fig 4E-G**).

### DNMT3B and DNMT3A are highly expressed in UPS and associated with a poor prognosis

Despite the *in vivo* toxicity of nanaomycin A-mediated DNMT3B inhibition, our studies of genetic *DNMT3B* depletion models (**Figs 2-3**, **EV Figs 1-2**) indicated that this enzyme nevertheless appears to be a promising molecular vulnerability in UPS. Therefore, to determine the extent to which continued development of pharmacologic DNMT3B inhibitors would have clinical utility, we characterized DNMT3B expression patterns in UPS and normal mesenchymal tissues. Given that DNMT3A depletion also showed anti-neoplastic activity *in vivo*, albeit to a lesser extent than that of DNMT3B, we also considered DNMT3A in this analysis in the event development of DNMT3B-specific inhibitors is ultimately not feasible due to unfavorable pharmacokinetic/pharmacodynamic profiles or toxicity concerns. Analysis of samples in the UPenn cohort and Detwiller sarcoma dataset demonstrated strong upregulation of *DNMT3B* gene expression in UPS/(M)FS samples relative to normal muscle and connective tissue (**Fig 5A-B**). *DNMT3A* gene expression was generally more stable than that of *DNMT3B*, particularly in the UPenn cohort, but was increased in a subset of UPS/FS specimens in the Detwiller dataset (**Fig 5A-B**). Similar trends were observed in canine samples from the Penn Vet cohort (**Fig 5C**). We also performed immunohistochemistry (IHC) for DNMT3B and DNMT3A on human sarcoma tissue microarrays (TMAs) containing UPS and normal skeletal muscle specimens (n = 11 and n = 5, respectively; **EV Fig 5A**; **Appendix Table S5**). These samples exhibited prominent nuclear immunoreactivity with relatively scant cytoplasmic staining (**Fig 5D**). Quantification of nuclear DNMT3B staining demonstrated significant upregulation in UPS compared to skeletal muscle tissue (**Fig 5E**, **EV Fig 5B**). Interestingly, in contrast to our gene expression results, nuclear DNMT3A expression was also significantly increased in tumors relative to normal muscle, suggesting that DNMT3A levels are subject to post-transcriptional regulation in UPS cells (**Fig 5E**, **EV Fig 5B**). Ultimately, greater than 70% of nuclei in all tumors examined were positive for at least one DNMT3 isoform. Consistent with this observation, DNMT3B and DNMT3A nuclear percent positivity were moderately correlated in UPS tumors (Pearson r = 0.5001; **Fig 5F**) according to conventional benchmarks for interpretation of correlation coefficients (Schober *et al*, 2018). In contrast, nuclear DNMT3B and DNMT3A positivity in normal skeletal muscle was more heterogeneous but more strongly correlated (Pearson r = 0.9302; **Fig 5G**).

**Fig 5.**
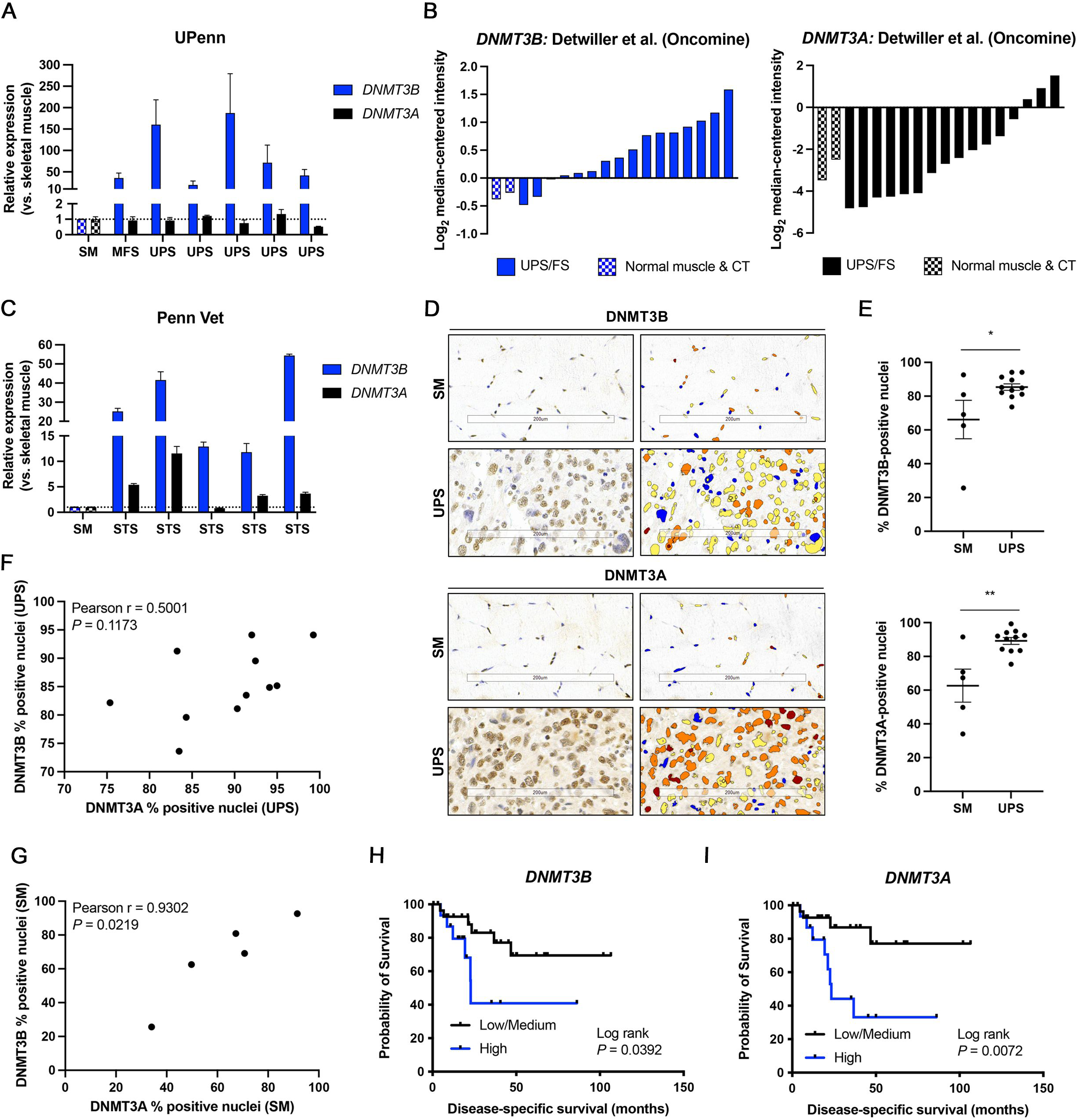
DNMT3B is overexpressed in human and canine UPS and associated with a poor prognosis. **A.** *DNMT3B* and *DNMT3A* gene expression in human UPS and MFS tumors relative to normal skeletal muscle (SM) tissue (UPenn cohort). Two cDNA samples per specimen. Mean ± SD. **B.** *DNMT3B* and *DNMT3A* gene expression in UPS and FS tissue specimens from the Detwiller et al. sarcoma dataset (Oncomine). CT = connective tissue. **C.** *DNMT3B* and *DNMT3A* gene expression in canine STS samples relative to normal skeletal muscle (SM) tissue (Penn Vet cohort). Mean ± SD. **D.** Representative images of DNMT3B and DNMT3A IHC staining and algorithm-based detection of nuclear immunoreactivity in normal skeletal muscle (SM) and UPS specimens (sarcoma tissue microarray; TMA). Red, orange, yellow, and blue represent 3+, 2+, 1+ and 0+ (negative) staining intensity, respectively. Scale bar = 200 µM. Image brightness and contrast have been enhanced for presentation purposes. **E.** Quantification of DNMT3B and DNMT3A percent positive nuclei in normal skeletal muscle (SM) and UPS specimens (sarcoma TMA). Two-tailed unpaired t-test. \**P* < 0.05, \*\**P* < 0.01. Mean ± SEM. **F.** Pearson’s correlation between the percentage of DNMT3B- and DNMT3A-positive nuclei in UPS specimens from panels **D-E**. Shapiro-Wilk W value = 0.9553. **G.** Pearson’s correlation between the percentage of DNMTB- and DNMT3A-positive nuclei in normal skeletal muscle (SM) specimens from panels **D-E**. Shapiro-Wilk W value = 0.9282. **H-I.** Disease-specific survival of human UPS patients in TCGA Sarcoma (TCGA-SARC) dataset stratified by tumor gene expression levels of *DNMT3B* (**H**) and *DNMT3A* (**I**). Each tertile (low, medium, and high) represents one-third of patients. Log-rank test. n = 44 patients.

Finally, we queried the Cancer Genome Atlas Sarcoma (TCGA-SARC) dataset to evaluate associations between the expression of DNMT-encoding genes and human UPS patient survival (n = 44). Consistent with our findings in UPS cell lines, expression of *DNMT1* was not associated with any survival endpoint (**EV Fig 5C-E**). However, high levels of *DNMT3B* and *DNMT3A* strongly tracked with reduced disease-specific (**Fig 5H-I**), overall (**EV Fig 5F-G**), and disease-free survival (**EV Fig 5H-I**). Taken together, these findings demonstrate that DNMT3B and DNMT3A are critical factors in UPS patient survival, and that modulation of their activity and/or expression may improve clinical outcomes.

## DISCUSSION

Comparative oncology is a powerful, often underutilized approach for elucidating fundamental mechanisms of tumor biology and developing novel therapeutic strategies (Rao *et al*., 2020). Due to the high prevalence of STS in pet dogs compared to human adults (∼15% vs. ∼1% of cases, respectively) (Katz *et al*., 2018; Rao *et al*., 2020), canine companion animals are particularly useful “models” for improving our understanding of these understudied cancers. Moreover, given several reports that STS subtypes, including UPS, are characterized by aberrant DNA methylation patterns (Cancer Genome Atlas Research Network. Electronic address & Cancer Genome Atlas Research, 2017; Kawaguchi *et al*., 2006; Koelsche *et al*., 2021; Merritt *et al*., 2018; Renner *et al*., 2013; Seidel *et al*., 2005; Seidel *et al*., 2007; Steele *et al*., 2019), careful examination of the underlying mechanisms and functional consequences of this epigenetic modification is critical. Herein, we adopted a comparative oncology approach consisting of human and canine patient-derived cell lines and tissue specimens, together with an orthotopic murine tumor model, to probe the function of DNMTs in UPS and evaluate their potential as anti-neoplastic targets. We discovered that DNMT3B genetic depletion in UPS cells inhibits proliferation *in vitro* and extends survival *in vivo*, but that an existing DNMT3B-targeting tool compound, nanaomycin A, elicits systemic toxicity, thereby precluding its clinical applicability.

Hypomethylating agents such as DAC and Azacitidine (Vidaza) received FDA approval for treatment of myelodysplastic syndrome nearly two decades ago. However, innate and acquired resistance to these drugs remains a significant clinical challenge (Fenaux *et al*, 2009; Jabbour *et al*, 2010; Prebet *et al*, 2011; Prebet *et al*, 2012; Qin *et al*, 2011). Herein, we showed that DAC resistance may also occur in the setting of UPS, potentially due to the downregulation of membrane-bound nucleoside transporters that facilitate intracellular drug uptake. Therefore, we assessed the anti-neoplastic efficacy of a potential alternative, nanaomycin A. To our knowledge, this work is the first to evaluate the therapeutic potential of nanaomycin A in any STS subtype. Consistent with other studies in neuroblastoma, melanoma, leukemia, and colon and lung carcinoma cell lines (Kuck *et al*., 2010a; Penter *et al*, 2015; Sztiller-Sikorska *et al*, 2014), we observed that clinically relevant, nanomolar doses of nanaomycin A potently inhibited UPS cell proliferation without inducing substantial cytotoxicity. However, its efficacy *in vivo* was extremely limited: although the highest evaluated dose almost completely ceased tumor growth in 2/10 animals, these same individuals experienced severe cachexia and were humanely euthanized. Body weights of the remaining mice in this group were also significantly lower than those of control mice. Notably, we are only aware of one other publication that evaluated the anti-neoplastic properties of nanaomycin A *in vivo*; however, data pertaining to tumor growth, animal survival, and systemic toxicity were not reported (Lai *et al*, 2019). Using computational molecular docking methods and biochemical assays, Kuck and colleagues (2010a; 2010b) previously demonstrated that nanaomycin A binds to the active site of DNMT3B, but not DNMT1. However, whether or not the compound also inhibits DNMT3A, which exhibits greater sequence homology to DNMT3B than does DNMT1, was not evaluated (Kuck *et al*., 2010a; Kuck *et al*., 2010b). Thus, we hypothesize that dual inhibition of DNMT3A and DNMT3B may at least partially underlie the observed systemic toxicity of nanaomycin A, particularly in light of reports that these paralogs can exhibit functional redundancy in some cell types (Challen *et al*, 2014; D’Antonio *et al*, 2012; Liao *et al*, 2015). Nevertheless, given the potency of nanaomycin A *in vitro*, we posit that development of novel pharmacologic DNMT3B-targeting strategies for UPS, and potentially other DNMT3B-overexpressing cancers, represents a promising avenue for future research.

Mechanistically, we found that genetic DNMT3B depletion blocks UPS cell proliferation by inhibiting DNA synthesis and causing cells to accumulate in G1-phase of the cell cycle. Similar findings have been reported in rhabdomyosarcoma, a pediatric skeletal muscle tumor (Megiorni *et al*, 2016), as well as in epithelial cancers such as pancreatic, bladder, and cholangiocarcinomas (Cao *et al*, 2021; Gao *et al*, 2013; Ying *et al*, 2020). In normal cells, G1-phase growth arrest is typically induced by p53 stabilization and transcription of its downstream targets (Agarwal *et al*, 1995). However, our work suggests that the growth arrest observed in DNMT3B-deficient UPS cells may be p53-independent, as KP230 cells in particular, like many human UPS tumors, are p53-null. In malignant contexts, p53-independent G1-phase growth arrest can occur via mechanisms such as suppression of c-Myc or downregulation of protein kinase C α and θ (Deeds *et al*, 2003; Jeong *et al*, 2010). Thus, evaluation of potential crosstalk between DNMT3B and these cell cycle regulators may provide further insight into the molecular susceptibilities of p53-deficient UPS cells.

Several challenges must be overcome before canine companion animals can be widely used as translational and clinical models of STS. First, functional studies in canine cells are currently limited by a paucity of cell lines and reagents (e.g., shRNAs, antibodies). Indeed, in the present study, we were unable to achieve consistent shRNA-mediated knockdown of canine *DNMT3B*, nor were we able to find any suitable shRNA constructs for depletion of *DNMT3A*. Of the experiments in which canine sarcoma cells were used, only one relevant cell line, STSA-1, was available. Thus, widespread commercialization of rigorously validated canine-specific reagents will facilitate broader incorporation of canine models into functional oncology and drug discovery research. In addition, as in the present study, detailed classification of canine STS is not routinely performed in veterinary medicine, largely due to poorly characterized associations among tumor subtype, patient prognosis, and therapeutic response (Boerkamp *et al*., 2016; Seguin, 2017). When detailed subtyping is performed, differences in nomenclature and classification conventions often lead to diagnostic inconsistencies between human and veterinary pathologists, even when cross-species tumor histologies are similar (Milovancev *et al*., 2015; Seguin, 2017). Therefore, development of a more reproducible STS classification system is paramount. Nevertheless, despite these challenges, our study underscores the promise of comparative oncology approaches. Our findings revealed strong parallels between human and canine cells/tissue with respect to multiple aspects of STS biology, including drug sensitivity/resistance patterns, as well as differences in gene expression between tumors and normal skeletal muscle. Ultimately, continued development of translational canine models in this area will provide valuable insight into the biology and treatment of this aggressive disease in both human and canine patients.

## MATERIALS AND METHODS

### Cell culture

HEK293T, HT-1080, and C2C12 cells were purchased from the American Type Culture Collection (ATCC, Manassas, VA). Human (HSMMs) and canine skeletal muscle myoblasts (CnSkMCs) were purchased from Lonza (Walkersville, MD) and Cell Applications, Inc. (San Diego, CA), respectively. STS-109 and STS-148 cells were derived from *TP53*-null UPS patient tumors and were a gift from Rebecca Gladdy, MD (University of Toronto). STSA-1 canine soft-tissue sarcoma cells were a gift from Molly Church, DVM, PhD (University of Pennsylvania School of Veterinary Medicine). Upon receipt of each cell line, multiple aliquots were frozen in liquid nitrogen within 10 days of initial resuscitation. For experimental use, aliquots were thawed and cultured for up to 20 passages (4-6 weeks) before being discarded. HEK293T, HT-1080, and KP230 cells were cultured in DMEM with 10% FBS, 1% L-glutamine, and 1% penicillin/streptomycin (P/S). STSA-1 cells were grown in DMEM with 15% FBS and 1% P/S. STS-109 and STS-148 cells were cultured in DMEM with 20% FBS, 1% L-glutamine, and 1% P/S. C2C12 cells were grown in DMEM with 20% FBS and 1% P/S. HSMMs and CnSkMCs were cultured in SkGM-2 (CC-3245, Lonza) and CnSkMC growth medium (Cn151-500, Cell Applications, Inc.) respectively. Sarcoma cell lines were not permitted to exceed 50% confluence during routine culture or experiments. Myoblast cell lines C2C12, HSMM, and CnSkMC were sub-cultured at 30-40% confluence. All cells were maintained in a humidified incubator at 37°C with 5% CO_2_ and confirmed to be negative for mycoplasma contamination.

### Oncomine and TCGA data analysis

The publicly available Detwiller et al. sarcoma dataset was used to analyze *DNMT3A* and *DNMT3B* expression levels in human UPS/FS and normal muscle specimens (access provided via Oncomine Research Premium Edition software, v4.5). For analysis of associations between *DNMT1*, *DNMT3A*, and *DNMT3B* gene expression and human UPS patient survival, the TCGA-SARC dataset was downloaded from the NIH Genomic Data Commons portal, imported into and normalized with *DESeq2* (Love *et al*, 2014) in R, and annotated with Ensembl BioMart. Clinical TCGA-SARC data (“TCGA, Cell 2017” dataset) were downloaded from cBioPortal on March 21, 2019; only cases included in the TCGA-SARC publication (Cancer Genome Atlas Research Network. Electronic address & Cancer Genome Atlas Research, 2017) were used in our analysis. Kaplan-Meier analyses were performed for disease-free, disease-specific, and overall patient survival.

### Lentiviral production and transduction

For shRNA-mediated knockdown studies, glycerol stocks of RNAi consortium (TRC) lentiviral vectors were purchased from Dharmacon (Lafayette, CO). A non-targeting control shRNA vector (sh:SCR) was purchased from Addgene (Watertown, MA). Plasmids were packaged with the third-generation lentiviral packaging system (VSV-G, pMDLg, and pRSV-REV) and expressed in HEK293T cells via transient transfection with FuGENE 6 (Promega, Madison, WI). Virus-containing supernatants were harvested after 24 and 48 hours, passed through 0.45 μm filters, mixed with PEG-8000 (Sigma-Aldrich, St. Louis, MO) at a 3:1 v/v ratio, and incubated overnight at 4°C. Supernatants were then centrifuged at 1500 x *g* for 30 minutes at 4°C and the virus-containing pellets were concentrated 80-fold in PBS. Aliquots were stored at −80°C and freeze-thawed a maximum of three times. For infection of sarcoma cell lines, cells were incubated overnight with lentiviral particles in the presence of 8.0 μg/mL polybrene (Sigma-Aldrich). Puromycin selection (3.0 μg/mL) was performed 48 hours after transduction and cells were harvested for analysis or used in further experiments after an additional 48 hours. Lentiviral constructs that exhibited the greatest knockdown efficiency in pilot analyses were used for all experiments: mouse *Dnmt3a*: TRCN0000039034 (sh:*Dnmt3a* #1), TRCN0000039036 (sh:*Dnmt3a* #2); mouse *Dnmt3b*: TRCN0000071069 (sh:*Dnmt3b* #1), TRCN0000071072 (sh:*Dnmt3b* #2); human *DNMT3A*: TRCN0000035755 (sh:*DNMT3A* #1), TRCN0000035757 (sh:*DNMT3A* #2); human *DNMT3B*: TRCN0000035685 (sh:*DNMT3B* #1), TRCN0000035686 (sh:*DNMT3B* #2). sh:*DNMT3B* #1 was not used in HT-1080 cells due to evidence of off-target effects (reduced proliferation without concomitant reductions in *DNMT3B* gene expression). To identify suitable shRNAs for use in canine sarcoma cells, all available mouse and human constructs were aligned to the canine *DNMT3A* or *DNMT3B* transcript (variant 1), as appropriate, using Standard Protein Blast (blastp). Acceptable shRNAs were defined as those that 1) possessed 100% query coverage in and 100% identity to the corresponding canine transcript, and 2) did not exhibit potential for off-target effects (defined as 100% query coverage in and 100% identity to a canine transcript other than the intended target). Based upon these criteria, human sh:*DNMT3B* #1 was the only shRNA suitable for use in canine cells (**Appendix Table S2**); however, we were unable to achieve consistent knockdown with this construct by qRT-PCR.

### Cell proliferation (growth curve) assays

For shRNA studies, cells were transduced with lentiviral particles as described above, trypsinized 96 hours post-infection, and seeded at equal densities in puromycin-containing culture medium for generation of growth curves. Cells were then counted using a hemocytometer with trypan blue exclusion every 2 days for 8 days (day 6, 8, 10, and 12 post-infection; STS-109 and STS-148 cells), or daily for 3 days (day 7, 8, and 9 post-infection; KP230, STSA-1, and HT-1080 cells). For siRNA studies, KP230 and HT-1080 cells were transfected with 50 nM mouse *Dnmt1* or human *DNMT1* ON-TARGETplus SMARTpool siRNAs with DharmaFECT reagent (Dharmacon) according to the manufacturer’s instructions. Cells were then counted every 24 hours for 3 days. For drug treatment studies, cells were treated daily with nanaomycin A (Apex BioTechnology, Houston, TX) for 3 days with enumeration every 24 hours (KP230, STSA-1, and HT-1080 cells), or for 8 days with enumeration every 48 hours (STS-109 and STS-148 cells). To prepare nanaomycin A for *in vitro* use, the drug was diluted in DMSO to a final concentration of 1 x 10^7^ nM and single use aliquots were prepared and stored at −80°C. All drug treatments were administered in fresh culture medium. Specific concentrations used for each cell line are indicated in the text.

### Orthotopic tumor murine model

Animal studies were performed in accordance with NIH guidelines and were approved by the University of Pennsylvania School of Medicine Animal Care and Use Committee. For *in vivo* knockdown studies, 6-week-old female nu/nu mice (strain code: 002019, The Jackson Laboratory, Bar Harbor, ME) were randomized to receive bilateral orthoptic injections (into the gastrocnemius muscles) of 1 x 10^5^ KP230 cells expressing a scrambled control (sh:SCR), *Dnmt3a*-(sh:*Dnmt3a* #1), or *Dnmt3b*-(sh:*Dnmt3b* #2) targeting shRNA lentivirus. Each mouse received bilateral injections of the same cell type to reduce animal use (n = 5 mice or 10 tumors/group). Tumors were measured with electronic calipers every 24-48 hours and tumor volume was calculated using the formula (ab^2^)π/6 where “a” and “b” represent the longest and shortest dimensions, respectively. Animals were euthanized when the first tumor on each mouse reached maximum allowed volume (1500 mm^3^).

For *in vivo* nanaomycin A treatment, 6-week-old female nu/nu mice (strain code: 002019, The Jackson Laboratory) received unilateral orthotopic injections of 1 x 10^5^ KP230 cells into the gastrocnemius muscle and were randomized to one of three groups: 0 mg/kg, 7.5 mg/kg, or 15 mg/kg nanaomycin A (n = 10 mice/group). Nanaomycin A treatment for each mouse began when its respective tumor became palpable, and the drug was administered intraperitoneally (i.p.; 100 μL injection volume) every 48 hours for up to 3 weeks or until tumors reached maximum allowed volume (1500 mm^3^). Tumor volume was measured and calculated as described above, and animal body weights were tracked to assess the impact of nanaomycin A treatment on overall health. To prepare nanaomycin A for *in vivo* delivery, the compound was reconstituted in DMSO to a final concentration of 5 x 10^7^ nM, and single-use aliquots were made by dissolving the appropriate volume of drug in an aqueous solution of 45% PEG-400 (Sigma-Aldrich); in each 100 μL injection, the final concentration of DMSO was 30% (v/v) for all groups. Aliquots were stored at −80°C until use.

### qRT-PCR

Total RNA was isolated from cells using the RNeasy Mini Kit (Qiagen, Germantown, MD) and from tissue (∼50-100 mg cut over dry ice) using TRIzol reagent (Thermo Fisher Scientific, Waltham, MA). Reverse transcription of mRNA (up to 2 μg/reaction) was performed using the Applied Biosystems High-Capacity RNA-to-cDNA kit (Thermo Fisher Scientific). qRT-PCR (20 ng cDNA/reaction) was performed using TaqMan probes (Thermo Fisher Scientific) on an Applied Biosystems ViiA7 instrument. All TaqMan probes were “best coverage” assays except for canine *DNMT3B* which was made with the TaqMan custom design tool (Custom Plus TaqMan RNA assay ID ARAADW9). Hypoxanthine phosphoribosyltransferase, succinate dehydrogenase subunit A, and/or beta-2 microglobulin were used as endogenous controls.

### RNA-sequencing and bioinformatics

Human STS-109 cells were transduced with *DNMT3A*- or *DNMT3B*-targeting shRNA lentiviruses (2 independent shRNAs per target) or a non-targeting shRNA control (sh:SCR) as described above, and lysed in preparation for total RNA isolation 96 hours after infection. RNA was extracted using the RNeasy Mini Kit (Qiagen) and further purified with the RNA Clean and Concentrator Kit (Zymo Research, Irvine, CA). Samples were quality-checked using the Agilent RNA 6000 Nano Kit and Agilent 2100 BioAnalyzer (Santa Clara, CA); all samples had RNA integrity values of >8.5. Poly(A) mRNA enrichment and library preparation were performed using the NEBNext Poly(A) mRNA Magnetic Isolation Module, NEBNext Ultra II RNA Library Prep Kit (New England BioLabs, Ipswitch, MA) with SPRIselect Beads (Beckman Coulter, Brea, CA), and NEBNext Multiplex Oligos for Illumina (index primers sets 1 and 2) according to the manufacturer’s instructions. Library sizes were checked with the Agilent DNA 1000 Kit, and library concentrations were determined with Qubit dsDNA assays (Thermo Fisher Scientific). Equimolar amounts of each library were pooled, and pools were diluted to a final concentration of 1.8 pM in HT1 Hybridization Buffer (Illumina, San Diego, CA) and sequenced on an Illumina NextSeq 500 instrument using the NextSeq 500/550 75-cycle High-Output Kit v2.5.

Illumina .bcl files were converted to FASTQ files using the Illumina *bcl2fastq* command line program. Salmon (Patro *et al*, 2017) was used to count raw expression data against the human transcriptome as defined in Gencode v32. Transcript-level data normalization and differential expression analysis (sh:*DNMT3A* vs. sh:SCR; sh:*DNMT3B* vs. sh:SCR) were performed with *DESeq2* (Love *et al*., 2014). To identify pathways that were differentially upregulated in sh:*DNMT3A* or sh:*DNMT3B* vs. control cells, we identified genes that were significantly upregulated (FDR-adjusted *P* < 0.05) in both shRNA comparisons for each target (i.e., genes upregulated in sh:*DNMT3A* #1 vs. sh:SCR <u>and</u> sh:*DNMT3A* #2 vs. sh:SCR). A similar process was carried out to identify genes that were significantly downregulated in both shRNA comparisons (i.e., genes downregulated in sh:*DNMT3A* #1 vs. sh:SCR <u>and</u> sh:*DNMT3A* #2 vs. sh:SCR). Additionally, to identify genes that were regulated by both *DNMT3A* and *DNMT3B*, we identified genes that were either up- or down-regulated in all 4 possible comparisons (i.e., genes significantly upregulated in sh:*DNMT3A* #1 vs. sh:SCR, sh:*DNMT3A* #2 vs. sh:SCR, sh:*DNMT3B* #1 vs. sh:SCR, <u>and</u> sh:*DNMT3B* #2 vs. sh:SCR). Pathway analyses on these “merged” gene lists were performed with Metascape (Zhou *et al*, 2019).

### Immunoblotting

Cells were lysed in 1x RIPA buffer supplemented with 1x protease and phosphatase inhibitors, and lysate concentrations were determined with the Pierce BCA Protein Assay Kit (Thermo Fisher Scientific). Denatured proteins were separated by electrophoresis on 8%, 10%, or 4-15% gradient sodium dodecyl sulfate polyacrylamide gels, transferred to 0.2 μm PVDF membranes using the Bio-Rad Trans-Blot Turbo System (Hercules, CA), blocked for 30-60 minutes in 5% non-fat dry milk, and probed with the following antibodies overnight at 4°C: rabbit anti-DNMT3A (0.2 μg/mL [1:500]; HPA026588, Atlas Antibodies, Stockholm, Sweden), rabbit anti-DNMT3B (0.4 μg/mL [1:500]; HPA001595, Atlas Antibodies), rabbit anti-Dnmt3b (1 μg/mL [1:1000]; NB300-516, Novus Biologicals, Centennial, CO), and rabbit anti-GAPDH (1:1000; #2118, Cell Signaling Technology, Danvers, MA). Membranes were then probed with horseradish peroxidase-conjugated anti-rabbit secondary antibodies (1:2500; #7074, Cell Signaling Technology) for 1 hour at room temperature. Chemiluminescent detection was performed with Western Lightning Plus-ECL reagent (PerkinElmer, Waltham, MA).

### Immunohistochemistry and digital histopathology

Human sarcoma TMAs were purchased from US Biomax, Inc. (SO801a, Derwood, MA). Murine tumors were formalin-fixed, paraffin-embedded, sectioned at 5 μm, and stained with hematoxylin and eosin according to standard protocols. Immunohistochemistry was performed on a Bond RXm autostainer instrument with the Bond Polymer Refine Detection kit (Leica Biosystems, Buffalo Grove, IL) using the following antibodies and epitope retrieval (ER) buffers: rabbit anti-DNMT3A (2 μg/mL [1:50], 30 minutes with ER1; HPA026588, Atlas Antibodies), rabbit anti-DNMT3B (0.4 μg/mL [1:500], 30 minutes with ER1; HPA001595, Atlas Antibodies), and rabbit anti-p62 (1:2000, 30 minutes with ER1; PM045, MBL International, Woburn, MA). Slides were scanned into digital images at a magnification of 20x using the Aperio VERSA 200 platform (Leica Biosystems). Images were annotated to exclude areas with poor resolution or that contained histologically normal cancer-adjacent tissue, folded tissue, debris, or staining/scanning artifacts. Analysis was performed with Aperio ImageScope software (Leica Biosystems) as follows: To assess nuclear DNMT3A and DNMT3B staining in tumor tissue, the “nuclear V9” algorithm was tuned to UPS cores from the human sarcoma TMA. To assess nuclear DNMT3A and DNMT3B staining in normal skeletal muscle tissue, skeletal muscle cores from the human sarcoma TMA were subjected to tissue composition analysis with the Aperio Genie Classifier. Use of this classifier was necessary to prevent detection/quantification of non-specific hematoxylin staining that was present in normal skeletal muscle cores but not in UPS cores. To train the composition estimator, normal skeletal muscle slides from an in-house tissue repository were randomly selected and manually annotated for skeletal muscle nuclear area, “other” tissue area, and glass. Following partitioning of each component, the same “nuclear V9” algorithms developed for assessment of DNMT3A and DNMT3B IHC staining in UPS cores were then applied to the “skeletal muscle nuclei” class of the normal skeletal muscle cores. Finally, the “positive pixel count” algorithm was used to quantify p62 staining in murine tumor tissue. A p62 IHC score was calculated as follows: (3 × % strong positive pixels) + (2 × % moderately positive pixels) + (1 × % weak positive pixels).

### TUNEL assay and immunofluorescence analysis

Murine tumor sections were deparaffinized and rehydrated according to standard protocols and microwave-irradiated with 10 mM sodium citrate buffer, pH 6.0, for 10 minutes. Detection of DNA fragmentation was performed with the *In Situ* Cell Death Detection Kit, Fluorescein (Roche, Indianapolis, IN) according to the manufacturer’s instructions. Hoechst 33342 (1 μg/mL) was used as a nuclear counterstain and coverslips were mounted with ProLong Diamond Antifade Mountant (Thermo Fisher Scientific). Images (5 fields per tumor section) were acquired with a Nikon Eclipse Ni microscope and Nikon NES Elements software with the pixel saturation indicator set to “on”. Areas with high levels of autofluorescence were avoided. Analysis of nuclear TUNEL staining was performed with Fiji: Watershed analysis of DAPI channel images (8-bit) was performed to “separate” nuclei that appeared to be touching. Nuclei were then converted to regions of interest (ROIs) that were “applied” to the corresponding TUNEL/fluorescein channel image (8-bit format). Staining intensity in these nuclear ROIs was then calculated as follows: integrated density normalized to number of nuclei.

### Cell cycle analysis and flow cytometry

KP230, STSA-1, and HT-1080 cells were transduced with lentiviral particles as described above or treated with nanaomycin A every 24 hours for 3 days; specific concentrations are noted in the text. Cells were pulsed-labelled with 10 μM EdU (Click-iT EdU Alexa Fluor 488 Flow Cytometry Assay Kit, Thermo Fisher Scientific) in fresh culture medium for 1 hour, harvested via trypsinization, and fixed in ice-cold ethanol for at least 2 hours. Detection of DNA synthesis was performed according to the manufacturer’s instructions, after which cells were stained with 1 μg/mL Hoechst 33342 at a concentration of 1 x 10^6^ cells/mL for evaluation of DNA content. Flow cytometric data were collected on an LSR II instrument with FACSDiva software (BD Biosciences, Franklin Lakes, NJ). The instrument was run on the lowest flow-rate setting (∼<200 events/second) and at least 20,000 singlet events were captured for each sample. Analysis of cell cycle distribution was performed with FlowJo software (BD Biosciences) in which compensation and gating were guided by single-color controls (equivalent to fluorescence-minus-one controls in this experiment). For all cell lines, sub-G1 cells were gated out because events in this region may represent cell fragments rather than whole cells (Riccardi & Nicoletti, 2006). Additionally, for STSA-1 cells, G2/M-phase and polyploid cells were also gated out due to 1) the inability of our protocol to distinguish diploid G2/M cells from tetraploid G1 cells), and 2) the highly variable proportion of polyploid cells in replicate experiments at baseline.

### Myoblast differentiation studies

To induce myoblast differentiation, C2C12, CnSkMC, and HSMM on gelatin-coated plates (Gelatin-based coating solution, Cell Biologics Inc., Chicago, IL) were allowed to recover overnight in growth medium, and were then switched to their respective differentiation medium (C2C12: sodium pyruvate-free DMEM + 2% horse serum + 1% P/S; CnSkMC: Canine Skeletal Muscle Differentiation Medium, Cell Applications, Inc.; HSMM: DMEM/F12 with 15 mM HEPES + 2% horse serum + 1 % P/S) for 6 days (C2C12) or 5 days (CnSkMC and HSMM). Cells were plated such that they would reach ∼80-90% confluence by the end of the differentiation period (C2C12: ∼2.1 x 10^4^ cells/cm^2^; CnSkMC: ∼1.0 x 10^4^ cells/cm^2^; HSMM: ∼1.0 x 10^4^ cells/cm^2^). In addition, proliferating cells grown overnight to ∼40-50% confluence (C2C12: ∼7800 cells/cm^2^; CnSkMC: ∼2.1 x 10^4^ cells/cm^2^; HSMM: ∼1.0 x 10^4^ cells/cm^2^) were used in each experiment to confirm successful induction of differentiation (qRT-PCR-based analysis of species-specific skeletal muscle differentiation markers).

To assess the impact of nanaomycin A on myoblast viability, myoblasts were differentiated as described above in the absence of nanaomycin A, and subsequently treated with nanaomycin A every 24 hours for 3 days. Percent viability was determined every 24 hours using a hemocytometer with trypan blue exclusion. Accutase was used to dissociate differentiated HSMMs because it was less toxic to these cells than trypsin. To assess the impact of nanaomycin A on myoblast differentiation to myotubes, cells were differentiated as described above in the presence or absence of nanaomycin A (daily treatments; specific concentrations used for each cell line are indicated in the text). HSMM, which underwent nanaomycin A treatment withdrawal, were maintained in nanaomycin A-free differentiation medium for an additional 5 days after the final drug treatment. For all experiments, nanaomycin A was diluted in DMSO and treatments were administered in fresh culture medium.

### Availability of data and materials

RNA-seq data from this publication has been deposited into the NCBI Gene Expression Omnibus (GEO) under accession number PENDING. All other data and materials are available from the corresponding author upon reasonable request.

## ACKNOWLEDGEMENTS

The authors wish to acknowledge Rebecca Gladdy, MD, of the University of Toronto for providing STS-109 and STS-148 human UPS cells, and Molly Church, DVM, PhD, for providing STSA-1 cells. We also thank John Tobias, PhD, of the UPenn Molecular Profiling Facility for assistance with bioinformatics. This work was funded by The University of Pennsylvania Abramson Cancer Center, The Penn Sarcoma Program, Steps to Cure Sarcoma, R01CA229688, and T32HL007971. The UPenn Molecular Pathology and Imaging Core, which provided routine histology services, is supported by P30DK050306. The UPenn Cytomics and Cell Sorting Resource Lab is supported by the Abramson Cancer Center, the Department of Pathology and Laboratory Medicine, the Immune Health Institute, and the Parker Institute of the University of Pennsylvania.

## AUTHOR CONTRIBUTIONS

**Ashley M. Fuller:** Methodology, validation, formal analysis, investigation, data curation, visualization, writing – original draft, writing – review and editing

**Ann DeVine:** Investigation

**Ileana Murazzi:** Investigation

**Nicola J. Mason:** Resources, funding acquisition

**Kristy Weber:** Resources, funding acquisition

**T. S. Karin Eisinger:** Conceptualization, methodology, resources, writing – review and editing, study supervision, funding acquisition

## CONFLICT OF INTEREST

The authors declare no conflicts of interest.

**EV Fig 1. DNMT3A and DNMT3B, but not DNMT1, regulate UPS cell proliferation *in vitro*. A.** Growth curves of KP230 cells transfected with control or *Dnmt1*-targeting siRNAs. Two-way ANOVA with Sidak’s. Not significant. Mean ± SEM. n = 3. **B.** qRT-PCR data demonstrating *Dnmt1* knockdown levels in the KP230 cells shown in A. Two-tailed unpaired t-tests. \*\*\*\**P* < 0.0001. Mean ± SEM. n = 3. **C.** Growth curves of HT-1080 cells transfected with control or *DNMT1*-targeting siRNAs. Two-way ANOVA with Sidak’s. Not significant. Mean ± SEM. n = 3. **D.** qRT-PCR data demonstrating *DNMT1* knockdown levels in the HT-1080 cells shown in C. Two-way ANOVA with Sidak’s. Not significant. Mean ± SEM. n = 3. **E.** Growth curves of STS-148 cells transduced with control or *DNMT3A*-targeting shRNAs. Two-way ANOVA with Dunnett’s (vs. sh:SCR). \**P* < 0.05, \*\**P* < 0.01; asterisks at each time point indicate significance relative to sh:SCR. Mean ± SEM. n = 3. **F.** Representative qRT-PCR data demonstrating *DNMT3A* knockdown levels in STS-148 cells. One-way ANOVA with Dunnett’s (vs. sh:SCR). \*\*\*\**P* < 0.0001. Mean ± SEM. n = 3. **G.** Growth curves of HT-1080 cells transduced with control or *DNMT3A*-targeting shRNAs. Two-way ANOVA with Dunnett’s (vs. sh:SCR). \*\*\*\**P* < 0.0001; asterisks at each time point indicate significance relative to sh:SCR. Mean ± SEM. n = 3. **H.** Representative qRT-PCR data demonstrating *DNMT3A* knockdown levels in HT-1080 cells. One-way ANOVA with Dunnett’s (vs. sh:SCR). \*\*\*\**P* < 0.0001. Mean ± SEM. n = 4. **I.** Growth curves of STS-148 cells transduced with control or *DNMT3B*-targeting shRNAs. Two-way ANOVA with Dunnett’s (vs. sh:SCR). \**P* < 0.05, \*\**P* < 0.01, \*\*\*\**P* < 0.0001; asterisks at each time point indicate significance relative to sh:SCR. Mean ± SEM. n = 3. **J.** Representative qRT-PCR data demonstrating *DNMT3B* knockdown levels in STS-148 cells. One-way ANOVA with Dunnett’s (vs. sh:SCR). \*\**P* < 0.01, \*\*\*\**P* < 0.0001. Mean ± SEM. n = 3. **K.** Growth curves of HT-1080 cells transduced with a control or *DNMT3B*-targeting shRNA. Two-way ANOVA with Sidak’s. \**P* < 0.05, \*\*\*\**P* < 0.0001. Mean ± SEM. n = 4. **L.** Representative qRT-PCR data demonstrating *DNMT3B* knockdown levels in HT-1080 cells. Two-tailed unpaired t-test. \*\*\*\**P* < 0.0001. Mean ± SEM. n = 3. **M.** Gene expression levels of *Dnmt3a* and *Dnmt3b* in bulk KP230 orthotopic tumor specimens. One-way ANOVA with Dunnett’s (vs. sh:SCR). \**P* < 0.05. Mean ± SEM. n = 10 tumors (5 mice) per group; 1 tumor in the sh:*Dnmt3b* group did not form. **N.** Full orthotopic KP230 tumor progression curves in nude mice (corresponds to **Fig 2J**). Mean ± SEM. n = 10 tumors (5 mice) per group; 1 tumor in the sh:*Dnmt3b* group did not form. **O.** Volumes of orthotopic KP230 tumors that had not reached maximum allowed volume upon euthanasia. One-way ANOVA with Dunnett’s (vs. sh:SCR). \**P* < 0.05; asterisk represents the ANOVA summary statistic. Mean ± SEM.

**EV Fig 2. Mechanistic analysis of DNMT3B and DNMT3A function in UPS. A-B.** Schematic (**A**) and Metascape pathway analysis (**B**) of genes significantly up- or down-regulated in sh:*DNMT3A*-expressing vs. sh:SCR human STS-109 cells (FDR *P* < 0.05). n = 4. **C**. Schematic showing identification of genes significantly up- or down-regulated in both sh:*DNMT3A* and sh:*DNMT3B*-expressing vs. sh:SCR human STS-109 cells (FDR *P* < 0.05). n = 4. **D.** Normalized nuclear TUNEL staining index in control and sh:*Dnmt3b* orthotopic KP230 tumors from nude mice. Two-tailed unpaired t-test. Not significant. Mean ± SEM. n = 10 tumors (5 mice) per group; 1 tumor in the sh:*Dnmt3b* group did not form. **E.** Representative images of the data quantified in **D**. Micrograph brightness and contrast have been enhanced for presentation purposes. **F-G.** IHC-based p62 expression score (**F**) and percent positivity (**G**) in control and sh:*Dnmt3b* orthotopic KP230 tumors from nude mice. Two-tailed unpaired t-test. Not significant. Mean ± SEM. n = 10 tumors (5 mice) per group; 1 tumor in the sh:*Dnmt3b* group did not form. **H.** Representative images of p62 staining and algorithm-based detection of p62 immunoreactivity in the tumors quantified in **F-G**. Red, orange, yellow, and blue represent 3+, 2+, 1+ and 0+ (negative) staining intensity, respectively. Scale bar = 200 µM.

**EV Fig 3. Nanaomycin a impedes UPS cell proliferation. A-B.** Growth curves of HT-1080 (**A**) and STS-148 (**B**) cells treated daily with nanaomycin a. Two-way ANOVA with Tukey’s. \**P* < 0.05, \*\*\**P* < 0.001, \*\*\*\**P* < 0.0001; asterisks represent significance relative to 0 nM. For ease of visualization, <u>only</u> statistical comparisons vs. 0 nM are presented in the figure. Full statistical results for all possible comparisons are shown in **Appendix Table S4**. Mean ± SEM. n = 3. **C.** Median fluorescence intensity of Hoechst 33342-DNA staining in the G1- and G2/M-phase KP230 cells depicted in **D** and **E**. **D.** Representative flow cytometry plots showing the cell cycle phase distributions of the control and nanaomycin a-treated KP230 and STSA-1 cells depicted in **C** and **E**. **E.** Quantification of KP230 and STSA-1 cell cycle phase distributions shown in **C** and **D**. Not significant. Mean ± SEM. n = 3.

**EV Fig 4. Evaluation of nanaomycin a toxicity in untransformed skeletal muscle cells and *in vivo*. A.** Representative data showing differentiation induction in murine (C2C12), canine (CnSkMC), and human (HSMM) myoblasts. Two-tailed unpaired t-tests. \**P* < 0.05, \*\**P* < 0.01, \*\*\**P* < 0.001, \*\*\*\**P* < 0.0001. Mean ± SD. “p” and “d” prefixes indicate proliferating and differentiated cells, respectively. **B.** Expression of murine skeletal muscle lineage markers in C2C12 myoblasts exposed to nanaomycin a or vehicle control over the course of differentiation to myotubes. One-way ANOVA with Dunnett’s (vs. sh:SCR). \**P* < 0.05. Mean ± SEM. n = 3. **C.** Expression of canine skeletal muscle lineage markers in CnSkMC myoblasts exposed to nanaomycin a or vehicle control over the course of differentiation to myotubes. qRT-PCR data for 250 and 500 nM were not acquired due to extensive cell death (≥ 50% and ∼100%, respectively). Independent replicates are shown due to heterogeneous responses. Two-tailed unpaired t-test. \**P* < 0.05, \*\**P* < 0.01, \*\*\**P* < 0.001, \*\*\*\**P* < 0.0001. Mean ± SD. Left panel: n = 2; right panel: n = 1. **D.** Enumeration of viable C2C12, CnSkMC, and HSMM myotubes after daily exposure to nanaomycin a or vehicle control. Cells were allowed to differentiate normally and treated with nanaomycin a or vehicle control every 24 hours for 3 days after differentiation was complete. 2-way ANOVA with Dunnett’s (vs. 0 nM). \**P* < 0.05, \*\**P* < 0.01, \*\*\**P* < 0.001, \*\*\*\**P* < 0.0001. Mean ± SEM. n = 3 for all cell lines except CnSkMCs, where n = 2; one replicate is not shown because of rapid, uniform nanaomycin a-induced cell death that prevented cell enumeration. **E.** Survival of orthotopic KP230 tumor-bearing nude mice treated with nanaomycin a or vehicle control. Log-rank test with Bonferroni multiple comparisons correction (table). n = 10 mice per group. **F.** Orthotopic KP230 tumor progression curves of control- or nanaomycin a-treated nude mice. n = 10 mice per group. **G.** Body weights of orthotopic KP230 tumor-bearing nude mice immediately prior to euthanasia. Two mice in the 15 mg/kg group that were humanely euthanized prior to the study endpoint are not shown. One-way ANOVA with Dunnett’s (vs. 0 mg/kg). \**P* < 0.05. Mean ± SEM. n = 10 mice per group.

**EV Fig 5. Associations between DNA methyltransferase gene expression and UPS patient survival in TCGA-Sarcoma (TCGA-SARC) dataset (n = 44). A.** High-power view of sarcoma patient tissue microarray (TMA) highlighting cores from UPS (green) and normal skeletal muscle (pink) specimens. **B.** Quantification of DNMT3B and DNMT3A percent positive nuclei in normal skeletal muscle (SM) and UPS specimens, stratified by nuclear staining intensity (sarcoma TMA). Two-tailed unpaired t-test (UPS vs. SM for each intensity category). \**P* < 0.05, \*\**P* < 0.01. Mean ± SEM. **C-E.** Disease-specific **(C)**, overall **(D)**, and disease-free (**E**) survival stratified by tumor expression levels of *DNMT1*. For all panels, each tertile (low, medium, and high) represents one-third of UPS patients. **F-G**. Overall survival stratified by tumor gene expression levels of *DNMT3B* **(F)** and *DNMT3A* **(G)**. For all panels, each tertile (low, medium, and high) represents one-third of UPS patients. **H-I.** Disease-free survival stratified by tumor gene expression levels of *DNMT3B* **(H)** and *DNMT3A* **(I)**. For all panels, each tertile (low, medium, and high) represents one-third of UPS patients.

